# O-GlcNAc regulates gene expression by controlling detained intron splicing

**DOI:** 10.1101/2020.03.27.012781

**Authors:** Zhi-Wei Tan, George Fei, Joao A. Paulo, Stanislav Bellaousov, Sara E.S. Martin, Damien Y. Duveau, Craig J. Thomas, Steven P. Gygi, Paul L. Boutz, Suzanne Walker

## Abstract

Intron detention in precursor RNAs serves to regulate expression of a substantial fraction of genes in eukaryotic genomes. How detained intron (DI) splicing is controlled is poorly understood. Here we show that a ubiquitous post-translational modification called O-GlcNAc, which is thought to integrate signaling pathways as nutrient conditions fluctuate, controls detained intron splicing. Using specific inhibitors of the enzyme that installs O-GlcNAc (O-GlcNAc transferase, or OGT) and the enzyme that removes O-GlcNAc (O-GlcNAcase, or OGA), we first show that O-GlcNAc regulates splicing of the highly conserved detained introns in *OGT* and *OGA* to control mRNA abundance in order to buffer O-GlcNAc changes. We show that *OGT* and *OGA* represent two distinct paradigms for how DI splicing can control gene expression. We also show that when DI splicing of the O-GlcNAc-cycling genes fails to restore O-GlcNAc homeostasis, there is a global change in detained intron levels. Strikingly, almost all detained introns are spliced more efficiently when O-GlcNAc levels are low, yet other alternative splicing pathways change minimally. Our results demonstrate that O-GlcNAc controls detained intron splicing to tune system-wide gene expression, providing a means to couple nutrient conditions to the cell’s transcriptional regime.

## INTRODUCTION

O-GlcNAc transferase (OGT), a glycosyltransferase that catalyzes the post-translational addition of O-linked N-acetylglucosamine (O-GlcNAc) to serine and threonine side chains of proteins, is required for viability of all mammalian cells (1,2). Four key features distinguish OGT as a unique glycosyltransferase that is distinct from canonical eukaryotic glycosyltransferases, which act exclusively in the secretory pathway to assemble the oligosaccharides found on cell surface glycoproteins. First, OGT and its substrates are found in the nucleus and cytoplasm and not in compartments of the secretory pathway such as the endoplasmic reticulum and the Golgi body (3). Second, OGT has over a thousand protein substrates that belong to virtually all classes of proteins in the cell, including structural and trafficking proteins, transcription factors, epigenetic regulators, components of the ribosome, the proteasome, and the kinome (4-7). Third, the modifications installed by OGT are dynamic due to the presence of a dedicated glycosidase, O-GlcNAcase (OGA), which removes O-GlcNAc modifications (8). In mammals, OGT and OGA are solely responsible for dynamic cycling of O-GlcNAc modifications (9,10). Fourth, O-GlcNAc levels change with nutrient conditions and are especially responsive to glucose levels (11). Because the concentration of OGT’s substrate, UDP-GlcNAc, increases with nutrient abundance, OGT is proposed to transduce overall nutrient levels, as reflected in UDP-GlcNAc concentration, into a signaling response that affects myriad cellular pathways (12-14). These pathways include phosphorylation networks that control metabolism, cell cycle regulation, and transcriptional regulation (15).

Many signaling processes act at short time scales. Because compounds that rapidly and specifically reduce O-GlcNAc through inhibition of OGT were not available until recently, studies that interrogate OGT function have primarily focused on changes that occur long after an initial perturbation has been applied (∼24-48 h) (16,17). Much has been learned about OGT, but the long time frame of typical studies is a limitation: some of the observed pathway changes may be downstream of the initial changes, and important effects may have been missed due to compensatory changes that occur prior to interrogation.

We have developed a class of OGT inhibitors useful for interrogating acute cellular changes that occur when OGT function is disrupted (18,19). Here we probe system-wide effects of OGT inhibition at short time points. We initially asked how the phosphoproteome changes because numerous studies have shown changes in protein phosphorylation when O-GlcNAc levels are reduced, suggesting extensive crosstalk between phosphorylation and O-GlcNAc signaling pathways (20-22). Quantitative phosphoproteomics showed changes in a large number of RNA splicing factors upon acute inhibition. Deep sequencing of polyadenylated RNA to probe how O-GlcNAc affects splicing showed distinct effects on detained introns, a specific form of alternative splicing.

Most introns are removed co-transcriptionally—that is, they are completely spliced prior to transcriptional termination and polyadenylation. In contrast, detained introns (DIs) are individual introns that remain in otherwise completely spliced, polyadenylated (post-transcriptional) messages, causing them to be detained in the nucleus rather than exported to the cytoplasm for translation (23). Many human gene transcripts contain detained introns, and incomplete splicing of these transcripts serves to control levels and timing of expression of productive (protein-coding) mRNAs. Based primarily on analysis of *Clk1* and *Clk4*, it was initially proposed that detained intron transcripts serve as a nuclear reservoir of potentially productive pre-mRNA that can be spliced if needed to increase protein abundance rapidly; indeed, this mode of action was recently shown to be important for neuronal activity responses (24-26). However, evidence now shows that not all detained introns function in this way. For example, some detained intron transcripts represent a “dead-end” because they cannot be spliced to functional message. They undergo decay in the nucleus or are spliced to a non-functional transcript in response to an environmental signal (23,27-31). Regardless of how detained intron pathways function for any given gene, it is believed that detained intron splicing serves as a mechanism to fine tune mRNA abundance without changing basal levels of transcription. By fine tuning gene expression, detained intron splicing provides an important mechanism by which cells carry out cell state transitions or homeostatic responses (32). Consistent with this idea, detained introns are frequently found in genes expected to be sensitive to environmental changes, including genes important in metabolism, stress responses, and differentiation (23,27).

A central hypothesis is that O-GcNAc serves as a signal to integrate metabolism with cell state, but how it does so is not known (33). We found two major, temporally distinct responses to OGT inhibition. After a 2 hour inhibitor treatment, cells responded to O-GlcNAc perturbation with highly specific changes in *OGT* and *OGA* splicing pathways, and we have shown that the resultant changes in splicing inversely regulate their productive mRNA levels. These splicing changes alter OGT and OGA protein levels to buffer changes in O-GlcNAc. Second, after 6 hours of OGT inhibitor treatment, over 80% of detained introns decreased, a remarkable response suggesting a coordinately regulated program for cell state transition. We conclude that when the initial rapid buffering response does not return cells to O-GlcNAc homeostasis, there are widespread changes in mRNA levels for almost all genes subject to detained-intron splicing control. Because we did not alter nutrient levels in these studies, our studies establish that O-GlcNAc is the direct signal for a nutrient-dependent response that changes gene expression by altering splicing pathways.

## MATERIAL AND METHODS

### Cell lines and cell culture

HEK-293T cells were grown in Dulbecco’s Modified Eagle Medium (Gibco, USA) supplemented with 10% FBS and 1X Penicillin-Streptomycin solution (Corning) at 37°C in 5% CO_2_. HCT116 cells were grown in McCoy’s 5A medium (thermo Fisher Scientific) supplemented with 10% FBS and 1X Penicillin-Streptomycin solution (Corning) at 37°C in 5% CO_2_. MEF cells were grown in Dulbecco’s Modified Eagle Medium (Gibco, USA) supplemented with 10% FBS and 1X Penicillin-Streptomycin solution (Corning) at 37°C in 5% CO_2_. OSMI-2 was synthesized as previously described, cycloheximide (239765-1ML) and Thiamet-G (SML0244) was purchased from Sigma-Aldrich (19). For siRNA transfection, HEK293T cells were transfected with the corresponding siRNA following the manufacturer’s protocol. Cells were harvested 2 days after treatment. The list of siRNAs used is provided in Supplementary Data (Supplementary Table S2).

### Western blotting

Cells were prepared for western blotting in the following manner, and all the steps were conducted at 4°C. Cells, treated with inhibitors or DMSO in fresh media, (changed 3 h before treatment) were washed once with cold PBS, collected in cold PBS, centrifuged and lysed in RIPA buffer (25 mM Tris, pH 8.0, 1% NP40, 0.5% DOC, 0.1% SDS, 150 mM NaCl) supplemented with cOmplete™ protease inhibitor cocktail (Sigma), PhosSTOP™ (Sigma), and 50 µM thiamet-G (Sigma). After this, samples were loaded on an SDS-PAGE gel and transferred to nitrocellulose membrane for immunoblotting. Antibodies used in this study include: anti-OGT (24083S, CST), anti-O-GlcNAc (RL2, ab2739, Abcam) and anti-actin (ab49900, Abcam).

### Immunoprecipitation

Cells cultured in 100 mM plates were treated either with 10 μM OSMI-2 or DMSO for 0.5 h or 1.0 h. Cells were washed and collected as indicated in the Western blotting section. At the indicated time point, cells were lysed in IP buffer (1% NP-40, 50 mM Tris, 2 mM EDTA and 150 mM NaCl) supplemented with cOmplete™ protease inhibitor cocktail, PhosSTOP™, and 50 µM thiamet-G, and homogenized by 10 passes through a 21-guage needle, and protein concentration was determined with the Pierce BCA protein assay kit (ThermoFisher). Insoluble materials were removed by centrifugation at 21,000 g for 30 mins at 4°C. The supernatant was precleared by incubation with Dynabeads protein G magnetic beads and anti-mouse IgG (sc-2025, Santa Cruz) for 1 h at 4°C. Protein concentrations of precleared lysates were determined by BCA assay and normalized before adding a mixture of two O-GlcNAc antibodies (Rl2 and CTD110.6). Precleared lysates were then incubated overnight at 4°C. The next day, Dynabeads protein G magnetic beads were added and samples were incubated for 2 h at room temperature. Beads were washed thrice with IP buffer and eluted with 0.5 M NH_4_OH solution. Samples were vacuum centrifuged to dryness and redissolved in 8 M urea, 100 mM EPPS pH 8.5. Samples were reduced with 5 mM TCEP for 15 mins at room temperature in the dark, then alkylated with 14 mM iodoacetamide for 45 mins at room temperature in the dark. Samples were then diluted 10-fold with 100 mM EPPS pH 8.5. Proteins were then digested with 1 μg LysC (Wako) overnight at room temperature and then with 1 μg sequencing grade trypsin (Promega) for 6 hours at 37°C. The resulting peptide solutions were then labelled with TMT 10/11-plex reagents (ThermoFisher). These fractions were subsequently acidified with 1% formic acid, vacuum centrifuged to near dryness and desalted with C18 stagetips. Dried peptides were resuspended in 5 % acetonitrile/5% formic acid for LC-MS^3^ processing.

### RNA sequencing

Cells in fresh media (changed 3 h before treatment) were treated either with compounds or DMSO for 2.0 or 6.0 h. Cells were washed and collected as indicated in the western blotting section. At the indicated time point, total RNA from OSMI-2 /DMSO samples (DMSO in triplicates and OSMI-2 in duplicates) and TMG/DMSO samples (DMSO in triplicates and TMG in triplicates) were isolated using Trizol (ThermoFisher) or RNeasy Plus kit (Qiagen) respectively, according to manufacturer’s directions. In Trizol isolated total RNA, Turbo DNase (ThermoFisher) was added to remove residual DNA. Agilent Bioanalyzer was used to determine RNA quality and integrity before isolation of poly(A)^+^ RNA. A total of 2 μg of RNA was used for subsequent steps. The TruSeq Stranded mRNA library preparation kit (Illumina) was used according to manufacturer’s instructions for a poly(A)^+^-based mRNA enrichment. The fragmentated mRNA samples were subjected to cDNA synthesis and library generation using the TruSeq mRNA library preparation kit (Illumina). Sequencing was performed with a NextSeq sequencer (Illumina) for paired end reads of ∼75 bp. Control, OSMI-2, and TMG-treated RNA-seq data is available from the Sequence Read Archive (SRA; GSE138783).

### Bioinformatics and statistical analyses

RNA-seq analysis were performed as previously described (23,27). Briefly, for OSMI-2/DMSO or TMG/DMSO samples (3 biological replicates each), an average of ∼42 M (OSMI-2) reads or ∼31 M (TMG) reads per sample were generated with an average successful alignment of ∼96% (OSMI-2) or ∼91% (TMG) using Star Aligner (34). For gene expression analysis, reads were aligned to hg19 reference genome using RSEM, and EBseq was used to perform subsequent differential expression analysis (35,36). Detained introns were identified as previously described (23). Using reads from OSMI-2/DMSO, reads were mapped using Bowtie and filtered with Bedtools. DESeq was then used to normalize intronic read counts (37-42). Alternative and constitutive intron classifications were performed using custom Python scripts, and are agnostic with regard to existing annotations other than known gene boundaries. Annotated reads were then used as an input to generate an ‘exon part’ gtf that was compatible with DEXSeq (43). The same generated gtf from OSMI-2/DMSO samples is used for TMG/DMSO analysis. Reads were counted from mapped .bam files using the counting script included with DEXSeq to generate count tables for each exon part. Differential expression of the alternative splicing events and detained introns was then determined using standard DEXSeq analysis with a padj. <0.05 as the cutoff for significant changes. One of the OSMI-2 treated samples appeared to have a dosing error because it clustered with the DMSO samples, and based on the high reproducibility of the OSMI-2 treatments validated by RT-qPCR, we excluded that replicate from subsequent analysis.

### Phosphopeptide enrichment

Cells were grown in a 150 mm dish in fresh media (changed 3 h before treatment) and were treated either with OSMI-2 or DMSO for 0.5 or 2.0 h. Cells were washed and collected as indicated in the western blotting section. At the indicated time point, cells were lysed in lysis buffer (2 % SDS 50 mM Tris and 150 mM NaCl), sonicated (BioDisruptor) and protein concentration determined with the BCA assay. Samples were reduced with 5 mM DTT for 45 mins at 60°C, then alkylated with 14 mM iodoacetamide for 45 mins at room temperature in the dark. Then 1 mg of protein was precipitated using chloroform/methanol. Protein pellets were resuspended in 200 mM HEPES pH 8.5 to 1 mg/mL. Proteins were digested with LysC (Wako) (substrate:enzyme=100) overnight at 37°C and then with sequencing grade trypsin (Promega) (substrate:enzyme=100) for 6 hours at 37°C. Equal amounts of peptide samples were desalted on a Waters C18 solid phase extraction Sep-Pak. The resulting peptide solutions were resuspended in 2 M lactic acid 50% ACN to a concentration of 2 mg/ml. Titansphere TiO_2_ beads (Gl Sciences) were washed twice with 2 M lactic acid 50% ACN and added into the peptide mixture to enrich for phosphopeptides. Phosphopeptides were washed twice with 1% TFA in 50% ACN and eluted with elution buffer (50 mM KH_2_PO_4_ pH 10). Formic acid was then added to a final concentration of 4% to acidify the solution. Phosphopetides were desalted and then labelled with TMT 10/11-plex reagents (ThermoFisher) for subsequent quantitative proteomics. TMT-labeled peptide samples were fractionated via basic-pH reversed-phase (BPRP) HPLC to 96 fractions and then consolidated to 12 fractions. These fractions were subsequently acidified with 1% formic acid, vacuum centrifuged to near dryness and desalted with C18 stagetips (44). Dried peptides were resuspended in 5 % acetonitrile/5% formic acid for LC-MS^3^ processing.

### Mass spectrometry and LC-MS/MS measurement

Our mass spectrometry data were collected using an Orbitrap Fusion Lumos mass spectrometer (ThermoFisher Scientific, San Jose, CA) coupled to a Proxeon EASY-nLC 1200 liquid chromatography (LC) pump (ThermoFisher). Peptides were separated on a 100 μm inner diameter microcapillary column packed with 35 cm of Accucore C18 resin (2.6 μm, 150 Å, ThermoFisher). For each analysis, we loaded ∼2 μg onto the column.

Separation was in-line with the mass spectrometer and was performed using a 3 h gradient of 6 to 26% acetonitrile in 0.125% formic acid at a flow rate of ∼450 nL/min. Each analysis used a MS3-based TMT method, which has been shown to reduce ion interference compared to MS2 quantification (45,46). The scan sequence began with an MS1 spectrum (Orbitrap analysis; resolution 120,000; mass range 400−1400 *m/z*; automatic gain control (AGC) target 5 × 10^5^; maximum injection time 100 ms). Precursors for MS2/MS3 analysis were selected using a Top10 method. Following acquisition of each MS2 spectrum, we collected an MS3 spectrum using our recently described method in which multiple MS2 fragment ions were captured in the MS3 precursor population using isolation waveforms with multiple frequency notches.

Mass spectra were processed using a SEQUEST-based pipeline (47). Spectra were converted to mzXML using a modified version of ReAdW.exe. Database searching included all entries from the human UniProt database. This database was concatenated with one composed of all protein sequences in the reversed order. Searches were performed using a 50 ppm precursor ion tolerance for total protein level analysis. The product ion tolerance was set to 0.9 Da. These wide mass tolerance windows were chosen to maximize sensitivity in conjunction with Sequest searches and linear discriminant analysis (48). TMT tags on lysine residues and peptide N-termini and carbamidomethylation of cysteine residues were set as static modifications, while oxidation of methionine residues was set as a variable modification. Relative quantitation was performed as previously described (49).

### RT-qPCR analysis

Total RNA was isolated employing the RNeasy Plus kit (Qiagen) according to manufacturer’s protocol. Extracted RNA was reversed transcribed into cDNA using random hexamer priming and Superscript III (ThermoFisher) according to the protocol supplied by the manufacturer. For qPCR, cDNA templates were amplified, and the C_T_ values were quantified with PowerUp SYBR Green Master Mix (ThermoFisher), and normalized to actin as an internal control. Experiments were performed using a StepOnePlus Real-Time PCR System (Applied Biosystems). poly(A)-mRNA was isolated using Ambion® Poly(A)Purist™ MAG Kit (ThermoFisher) according to manufacturer’s protocol. Nuclear and cytoplasmic separation of RNA was performed with Thermo Scientific NE PER Nuclear and Cytoplasmic Extraction Kit (ThermoFisher) according to manufacturer’s protocol. RNA in either the nuclear pellets or cytoplasmic fractions was extracted with RNeasy Plus kit (Qiagen). Total RNA extracted per fraction was quantified and used to calculate the percentage of mRNA localization for the RT–qPCR quantification. The list of primers is provided in Supplementary Data (Supplementary Table S1).

## RESULTS

### Acute OGT inhibition changes splicing factor phosphorylation

We hypothesized that changes in phosphorylation would identify processes rapidly affected by acute OGT inhibition. OSMI-2 is a cell-permeable OGT inhibitor with a submicromolar K_d_ and a well-defined binding mode characterized by X-ray crystallography (Figure 1A) (19). This compound was shown previously to reduce global O-GlcNAc levels by 4 h of treatment in a number of cell lines, including HEK-293T cells (19,50). Because shorter treatment times were not examined, we first asked whether OSMI-2 reduced O-GlcNAc at earlier time points. We assessed O-GlcNAc levels after OSMI-2 treatment by immunoblotting with a commonly used antibody and observed a substantial decrease by 0.5 h (Figure 1B and Supplementary Figure S1A). We also performed a pulldown of O-GlcNAc-modified proteins after a 0.5 and 1 h incubation with OSMI-2 and observed a ∼30% and ∼50% depletion in protein abundance, respectively, in good agreement with the immunoblotting data (Supplementary Figure S1B). Although changes in global O-GlcNAc levels may underestimate the extent of changes in O-GlcNAc for some proteins, and overestimate them in others, we were satisfied that there was a sufficient perturbation by 0.5 h of inhibitor treatment to use this time point in further studies.

**Figure 1.**
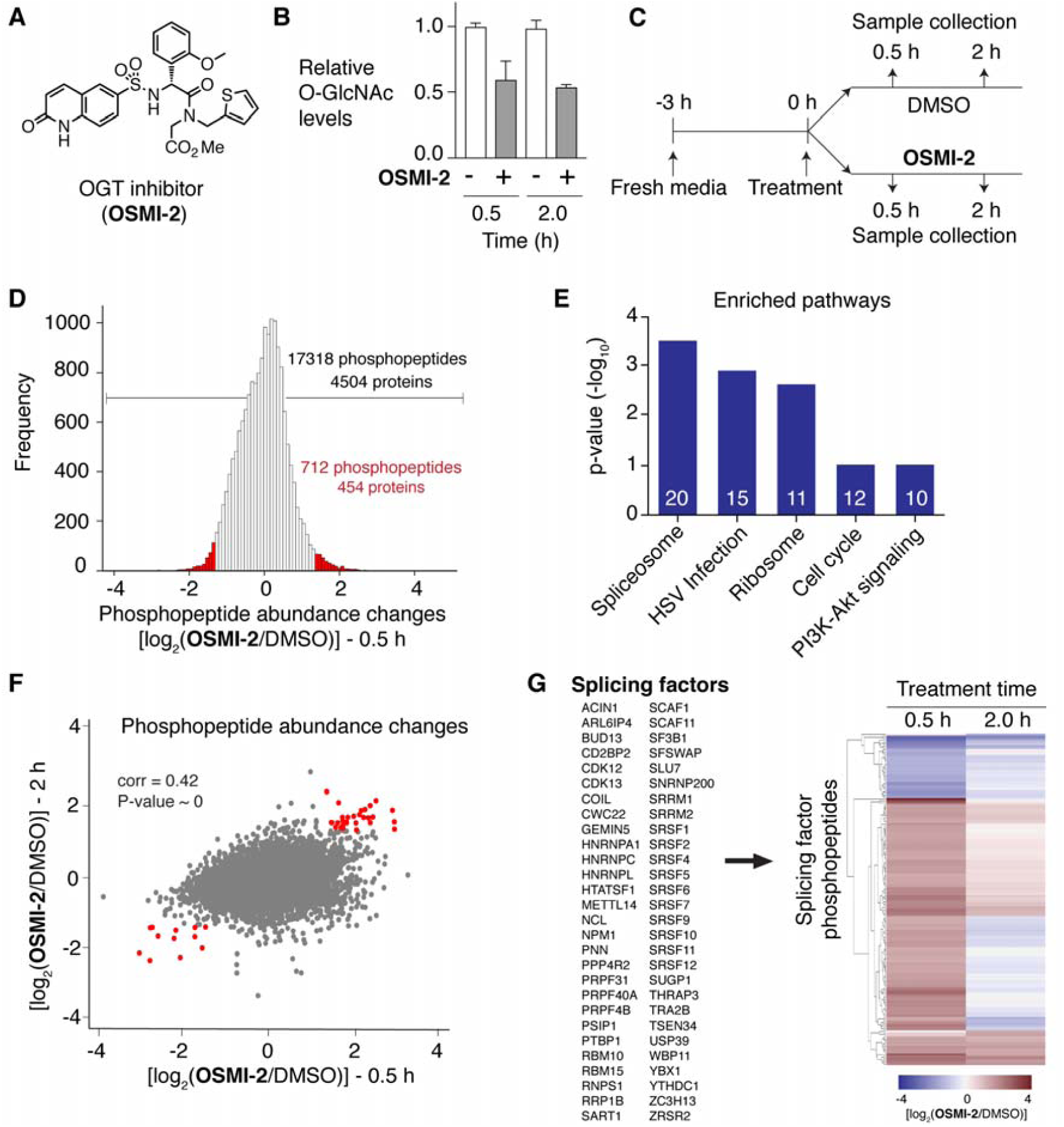
Phosphoproteome changes implicate OGT activity in RNA splicing. **(A**) Structure of OSMI-2. **(B**) Relative protein O-GlcNAc levels in HEK-293T cells show substantial changes by 0.5 h treatment with OSMI-2. (See also Supplementary Figure S1A) (**C**) Scheme depicting treatment conditions for quantitative phosphoproteomics. **(D)** Distribution of phosphopeptide abundances after 0.5 h treatment with OSMI-2, relative to DMSO control. Hits showed greater than a 2.5-fold change (2 STD from the median; shown in red). **(E)** KEGG pathway analysis of phosphoproteins that changed significantly by 30 minutes after OGT inhibitor treatment showed that the spliceosome was the most affected pathway. **(F)** Scatterplot of phosphopeptides showing changes at 0.5 h and 2 h treatment with OGT inhibitor (Pearson’s correlation coefficient = 0.42). Red dots represent hits meeting the cutoffs at both 0.5 h and 2 h. **(G)** Gene list and heatmap showing splicing factors with phosphosites that changed significantly at either 0.5 h or 2.0 h treatment with OGT inhibitor.

We asked how inhibiting OGT for 0.5 and 2 h with OSMI-2 affected global phosphorylation networks (Figure 1C). Using an isobaric-tagging approach (Tandem Mass Tagging, or TMT) for quantitative phosphoproteomics, we identified 17,318 phosphopeptides belonging to 4,504 proteins, of which 712 phosphopeptides showed at least a 2.5-fold change, corresponding to >2 standard deviations from the mean, at 0.5 h in the compound-treated sample compared with the DMSO control (Figure 1D and Supplementary Figure S2A). We analyzed the KEGG pathways that were enriched at 0.5 h and found some that were expected as well as some that were not (Figure 1E). For example, O-GlcNAc modifications have been shown previously to regulate the cell cycle and play a crucial role in PI3K-Akt signaling, and both of these pathways were enriched (51). Ribosomal components were also enriched and several studies have reported a link between O-GlcNAc and ribosome biogenesis and function (52-54). However, the top two enriched pathways, the spliceosome and HSV infection, have received little to no attention as pathways regulated by OGT. One previous study reported that OGT inhibition interferes with HSV replication through an undefined mechanism (55). Consistent with the fact that HSV replication relies on co-opting the splicing machinery of the host (56,57), when we examined the KEGG pathway genes associated with HSV infection, we found that half of them also belong to the spliceosome pathway. These results implicated RNA splicing as a process that changes rapidly upon OGT inhibition.

We asked whether the phosphorylation changes observed at 2 h recapitulated our observations at 0.5 h, highlighting RNA splicing as an affected pathway. Indeed, there was a good correlation between phosphorylation changes at 0.5 h and 2.0 h (Pearson’s correlation r = 0.42; *P* ∼0, Figure 1F). We identified a total of 55 splicing-related proteins that contained phosphopeptides showing at least a 2.5-fold change relative to the DMSO control at either 0.5 h (52 proteins), 2 h (14 proteins), or both time points (11 proteins) (Supplementary Figure S2B). Analysis of only those phosphopeptides that showed greater than 2.5-fold changes in the same direction at both time points identified RNA splicing as the only enriched process (Supplementary Figure S2C). The correlation analysis therefore confirmed splicing as a process to examine further.

### RNA-seq showed that acute OGT inhibition specifically affects detained introns in transcripts for *OGT* and *OGA*

Prior to this work, connections between RNA splicing and O-GlcNAc were limited. It was known that some splicing factors are O-GlcNAc-modified, but the functional significance of O-GlcNAc on splicing factors has not been examined (5). Previous studies showed that the *OGT* gene contains a detained intron, and it was known that levels of this intron respond to O-GlcNAc perturbation, indicating that *OGT* transcript splicing is subject to feedback regulation by O-GlcNAc (23,58). Despite the evidence that O-GlcNAc modification is involved in the splicing regulation of *OGT*, global transcriptome analysis has not been performed to assess whether detained introns in other genes are under O-GlcNAc control, nor have other splicing modalities been examined for O-GlcNAc-regulated changes.

To test whether any detained introns other than those of *OGT* are affected by acute OGT inhibition, we performed RNA-seq on poly(A)-selected RNA after 2 h or 6 h of treatment with OSMI-2. The 2 h timepoint was chosen because DI levels for *OGT* respond within this timeframe (58). Using our previously published bioinformatic methods, we identified DIs from the RNA-seq data and calculated the differential expression for each DI between control and inhibitor-treated samples to identify those that changed (23,43). At this early timepoint of 2 h after inhibitor treatment, we observed changes in DI splicing for only three genes: *OGT*, as expected; *OGA*, which encodes the glycosidase that together with OGT controls cellular O-GlcNAc levels; and *QTRTD1*, which encodes a subunit of tRNA-guanine transglycosylase, an enzyme that incorporates the hypermodified guanine analog queuosine into the wobble position of some tRNAs (Figure 2A). Although we did not follow-up on the latter observation, we point out that the queuosine tRNA modification is proposed to serve as a nutrient-controlled mechanism to fine-tune protein translation (59). Normalized RNA-sequencing read depths in the regions containing the DIs clearly exhibited increased read depth compared to other introns within the genes (Figure 2B). Upon OGT inhibition (OSMI-2), the read density mapping to the DIs in both genes decreased. Although introns typically exhibit poor sequence conservation, it was previously noted that the DI in OGT contains an ‘ultraconserved’ region (58), and indeed the vertebrate conservation tracks demonstrate that the DIs in both *OGT* and *OGA* are extensively embedded with highly phylogenetically conserved regions. The phylogenetic conservation of the *OGT* and *OGA* DIs, combined with quantitative global RNAseq analysis indicating these DIs decrease in response to OGT inhibition, suggests that a specific and conserved O-GlcNAc-responsive mechanism controls splicing of both the *OGT* and *OGA* transcripts to regulate protein abundance.

**Figure 2.**
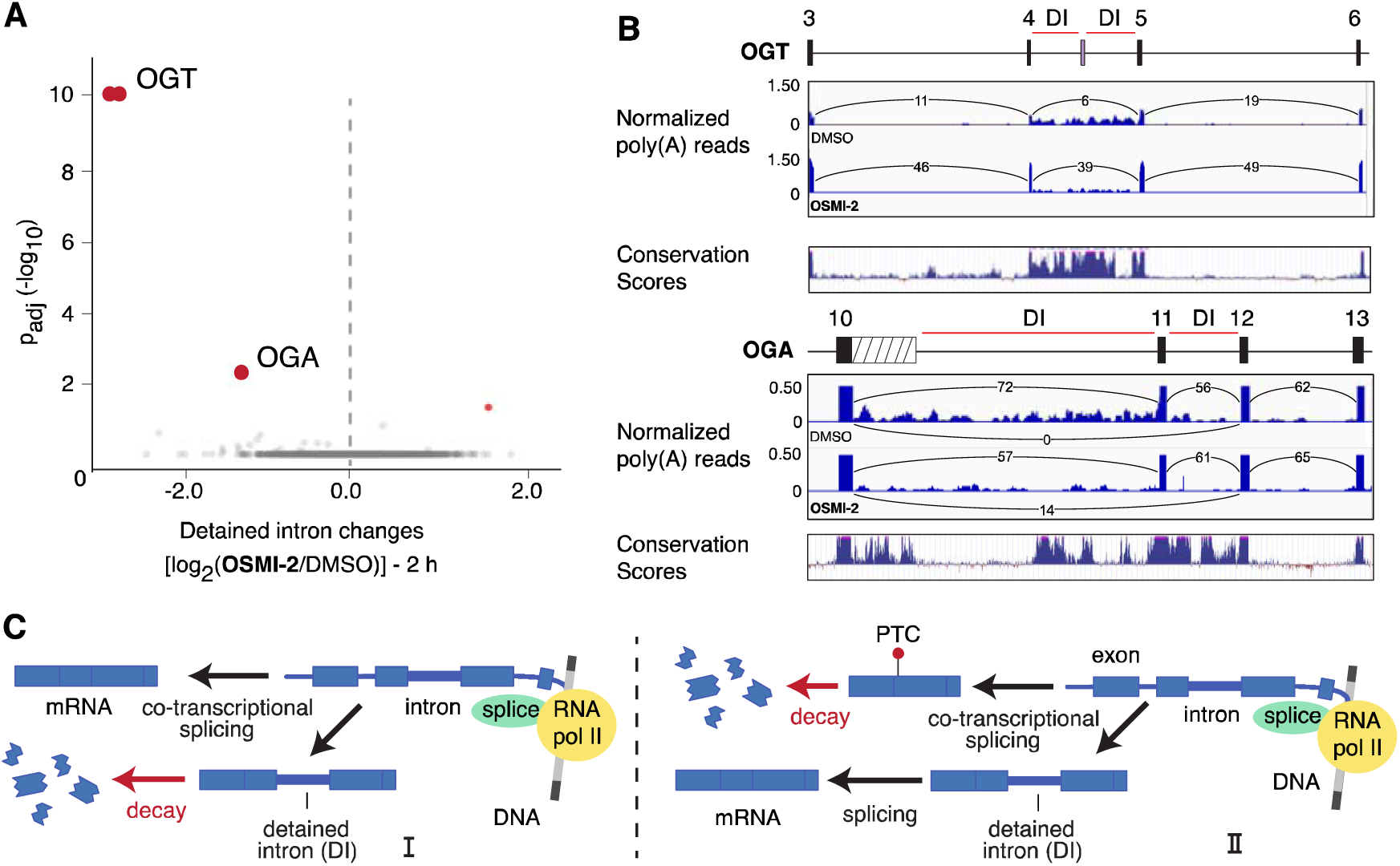
Acute OGT inhibition specifically regulates the highly conserved DIs of *OGT* and *OGA*. **(A)** Volcano plot showing fold changes (log_2_) in DIs after treatment with 10 μM OSMI-2 for 2 h. Inhibiting OGT specifically decreases DIs in *OGT* and *OGA*. The small red dot shows increased DI levels in *QTRTD1*. **(B)** Schematic of the *OGT* and *OGA* genes followed by total RNA-seq reads and vertebrate conservation track for the corresponding regions. Black boxes denote exons, and striped box denotes a short isoform of OGA produced by alternative polyadenylation. The top histograms show poly(A) reads for both DMSO and OSMI-2 treatment conditions normalized with total poly(A) reads per treatment. The Y-axis is the number of polyA reads normalized to read density and is given in terms of reads per million mapped reads. The number of junction spanning reads for relevant junctions are also given. The bottom histograms show the vertebrate conservation tracks obtained from the UCSC Genome Browser and indicate nucleotide-level conservation across 100 vertebrates, with pink caps indicating regions of 100% identity. **(C)** Scheme showing two different models for formation of productive mRNA through DI splicing. In pathway I, co-transcriptional splicing to make productive message competes with DI retention. In this model, the DI transcript is degraded. In pathway II, co-transcriptional splicing generates unproductive mRNA in competition with formation of a DI transcript that leads to productive mRNA.

Previous work identified the intron remaining in *OGT’s* transcripts as a *bona fide* DI because it caused the transcript to be detained in the nucleus (58). To confirm that the intron identified in *OGA* is also a DI resulting in nuclear detention of the transcript, we performed subcellular fractionation and used RT-qPCR to quantify transcript abundance in the nucleus and cytoplasm. We confirmed previous results for OGT, and found that OGA’s DI-containing transcript is also enriched in the nucleus compared to the fully spliced OGA transcript or to the transcript encoding actin (*ACTB*), which was used as a control (Supplementary Figure S3A). The DI splicing changes we observed at 2 h were remarkably selective, with *OGT* and *OGA* among the three genes showing changes. It was also initially surprising that DI levels decreased for both *OGT* and *OGA*; however, it is increasingly appreciated that DIs can control gene expression in different ways. For example, some studies have shown that when cellular conditions change, the fraction of nascent transcripts that undergo co-transcriptional splicing versus intron detention can be dynamically controlled to rapidly adjust protein levels (24,27,60). For some genes, co-transcriptional splicing evidently competes with intron detention that, in turn, leads to nuclear decay (Figure 2C, pathway I). For other genes, the detained-intron pathway leads to productive messages. In these cases, cellular signals may affect co-transcriptional splicing to favor the detained intron pathway and post-transcriptional splicing to productive message may also increase (Figure 2C, pathway II).

### O-GlcNAc-responsive DI splicing regulates *OGT* and *OGA* mRNA transcript abundance

We examined both the gene sequences and our RNA-seq data for clues to how the *OGT* and *OGA M*DIs may function. *OGT* was reported to contain a detained intron between exons 4 and 5, but closer inspection revealed that this region actually consists of two DIs flanking a small cassette exon (Figure 3A) (23,58). This same cassette exon was recently described as belonging to the class of ‘decoy exons’, exons upon which pre-spliceosome assembly stalls, resulting in two flanking DIs (61). This decoy exon is also termed a ‘poison’ exon due to the presence of a premature termination codon (PTC) within its sequence. Fully-spliced mRNA transcripts that contain poison exons undergo nonsense-mediated decay (NMD) upon export to the cytoplasm and translation (62). *OGA* also contains two conserved DIs that flank an exon, but in this case the exon (exon 11) is found in the productive, protein-coding *OGA* transcript. The RNA-seq data revealed an exon 11-skipped transcript containing a premature stop codon that was introduced by the reading-frame shift resulting from joining exon 10 and exon 12 (Figure 3B). Thus, *OGT* and *OGA* share a common structure comprising a central cassette exon flanked on both sides by DIs, with the splicing outcome of inclusion versus skipping producing opposite effects on the production of coding mRNA for each gene.

**Figure 3.**
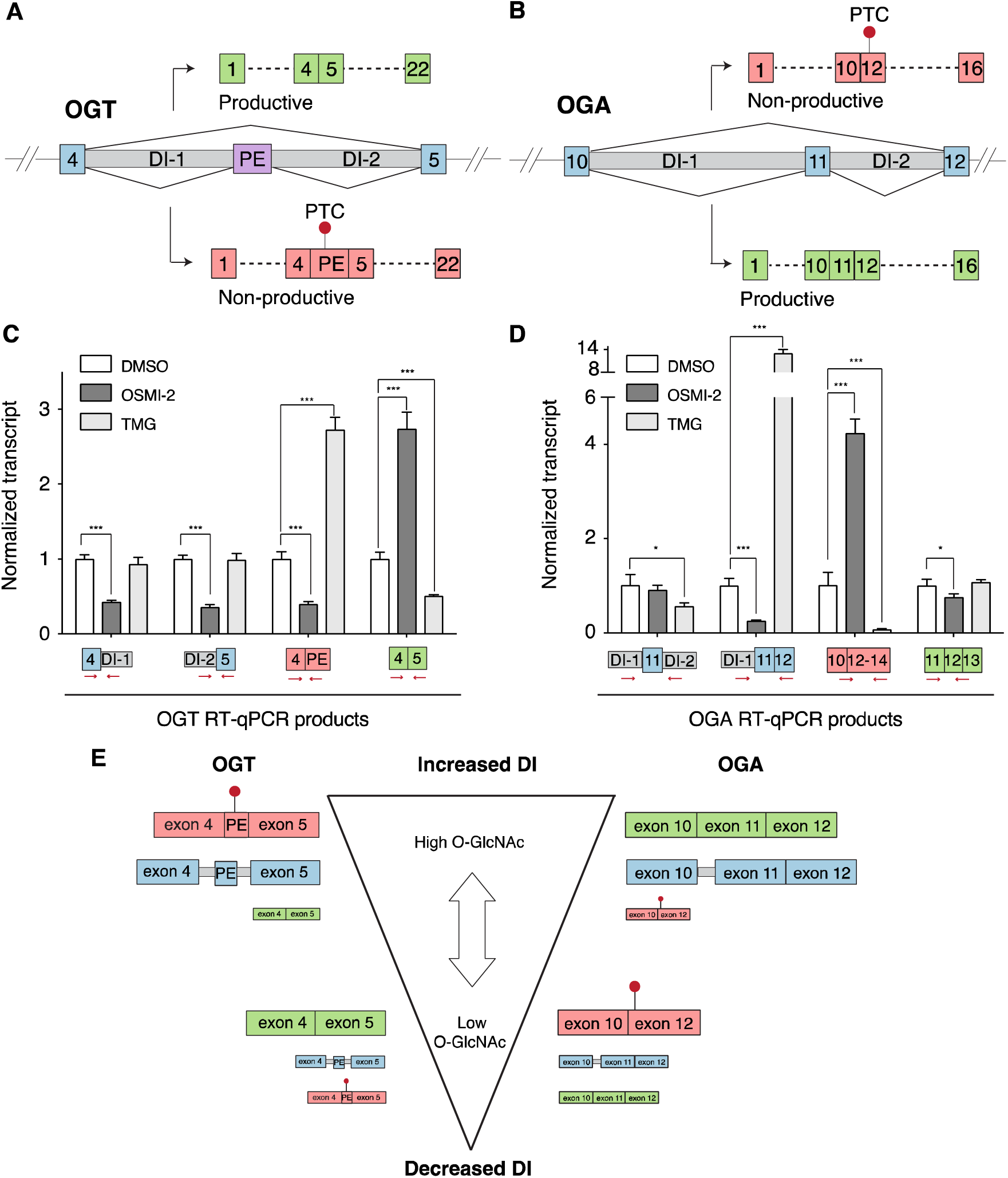
O-GlcNAc levels regulate *OGT* and *OGA* DI splicing so that exon inclusion/skipping results in opposite effects on productive message. **(A)** Splice junctions derived from RNA-seq for *OGT* and *OGA* show two predominant fully-spliced isoforms, a functional one for *OGT* **(A)** and a non-functional one for *OGA* **(B)**. HEK-293T cells were treated with DMSO (white bars), OGT inhibitor (10 μM OSMI-2, dark grey) or OGA inhibitor (5 μM TMG, light grey) for 2 h and RT-qPCR was used to measure levels of the indicated products for OGT **(C)** or OGA **(D)**, which report on the presence of DIs as well as skipped or included exons. Red arrows depict location of primers with respect to detected products. Products levels were determined relative to a housekeeping gene (actin) and were normalized to the DMSO control (n ≥ 3 biological replicates; mean ± s.d, \**P* ≤ 0.05, \*\*\**P* ≤ 0.001, two-tailed Student’s *t*-test). **(E)** Low O-GlcNAc favors the non-DI pathway to increase functional *OGT* mRNA and non-functional *OGA* mRNA. High O-GlcNAc favors formation of DIs, the splicing of which, increases non-functional *OGT* and functional *OGA*.

To confirm O-GlcNAc regulation of DI splicing for *OGT* and *OGA*, we designed PCR primers to quantify levels of the different transcripts and splice products following treatment with either the OGT inhibitor OSMI-2, or thiamet-G (TMG), a well-validated OGA inhibitor (64). With OGT inhibition, both *OGT* DIs decreased and levels of the productive (protein coding) mRNA isoform increased (Figure 3C). These changes corroborate northern blot data obtained previously for *OGT* DI splicing (58). With OGA inhibition, the spliced, productive mRNA transcript decreased, and the non-productive mRNA transcript containing the poison exon increased. The poison exon-containing product increased more than three-fold in the presence of the protein translation inhibitor cycloheximide. Nonsense-mediated decay requires translation in the cytoplasm so these results are consistent with the poison exon product being an NMD substrate. In contrast, the DI-containing transcript did not increase significantly in the presence of cycloheximide despite containing a PTC, consistent with this transcript being protected from NMD by nuclear sequestration (Supplementary Figure S3B). Taken together, our results show that the RNA processing pathways to productive versus non-productive *OGT* transcripts oppose each other in an O-GlcNAc-dependent manner, confirming that O-GlcNAc regulates *OGT* DI splicing. We also identify poison exon inclusion as an important mechanism for downregulating productive *OGT* mRNA expression when OGA is inhibited.

We next probed O-GlcNAc regulation of *OGA* splicing, using primers that reported on transcripts containing both DIs, or only DI-1, and on the exon 11-skipped (non-productive) mRNA as well as the protein-coding mRNA containing exon 11. Cycloheximide treatment showed a large increase in the exon 11-skipped product, consistent with this also being an NMD substrate. As with *OGT*, the *OGA* DI transcript did not increase when protein translation was inhibited (Supplementary Figure S3B). In the presence of OGT inhibitor, we observed a decrease in both DI-containing transcripts, but with a substantially larger decrease for transcripts containing only DI-1 (Figure 3D). We also observed a large increase in the exon 11-skipped transcript and a statistically significant decrease in productive transcript, although not to an extent commensurate with the increase in the exon-skipped product. However, we note that productive mRNAs comprise ∼80% of *OGA* transcripts under steady-state conditions and the mRNA half-life is relatively long (∼8.5 h) (65); both of these factors would dampen the impact of splicing changes on productive mRNA levels at short time points. Indeed, with longer treatment time (6 h), we observed a substantial decrease in the productive transcript of *OGA* (Supplementary Figure S4). When OGA was inhibited, we again observed large changes in the amount of transcript containing DI-1 and in the exon 11-skipped transcript, but they were in the opposite direction from those observed under OGT inhibition. These results indicate that splicing of *OGA*, similar to *OGT*, is regulated by O-GlcNAc.

From our RT-qPCR analysis of different transcripts under O-GlcNAc perturbation, we propose a working model for how O-GlcNAc affects *OGT* and *OGA* splicing pathways (Figure 3E). For this model, we assume that co-transcriptional splicing produces fully spliced mRNA in competition with the detained intron pathway, and that nascent transcripts are directed to one of the two outcomes in a proportion determined by O-GlcNAc levels. For both *OGT* and *OGA*, high O-GlcNAc results in increased transcript flux into the detained intron pathway. In both cases, intron detention appears to require commitment to the inclusion of the central exon, consistent with the ‘decoy exon’ hypothesis (61). For OGT, this results in the observed higher levels of the exon 4-poison exon spliced product in TMG-treated cells. For *OGA*, the detained intron pathway evidently results in functional product. We infer this based on the observation that when OGA is inhibited, there is a very large increase in a DI transcript that contains DI-1 but not DI-2. Because exons 11 and 12 have been joined, this transcript is a direct precursor to functional mRNA. Further, because the joining of exon 11 and 12 obviates the skipping of exon 11 through the removal of the 3’ splice site, formation of this transcript competes with the formation of the non-productive isoform. As a result of their inverted, symmetrical gene structures, high O-GlcNAc levels – through the single mechanism of increasing intron detention – lead to the downregulation of OGT by increasing splicing to the non-productive NMD isoform, while increasing productive OGA mRNA through exon 11 inclusion. The resulting enzymatic activity would be directed toward removal of O-GlcNAc modifications to restore homeostasis.

When cellular O-GlcNAc levels are low, the balance shifts toward increased co-transcriptional splicing and away from intron detention. In both loci, increased co-transcriptional splicing appears to promote reduced inclusion of the central DI-flanked exon. In the case of *OGT*, low O-GlcNAc results in more functional mRNA by promoting the skipping of the poison exon, while in the case of *OGA*, low O-GlcNAc enhances splicing of the frame-shifted NMD substrate (Figure 3E). The rapid, reciprocal, O-GlcNAc-controlled changes we observed in *OGT* and *OGA* splicing pathways provide an explanation for a well-known observation, namely, that OGT and OGA levels change in opposite directions when either protein is blocked (18,19,50,66). These O-GlcNAc-regulated splicing changes provide an elegant mechanism for how a single signal can be read out to produce opposing gene expression outcomes through the structural configuration of exons and introns, enabling OGT and OGA levels to be precisely tuned to buffer nutrient fluctuations. In addition to the unusual degree of phylogenetic sequence conservation of these introns, we found further evidence that this mechanism is conserved by verifying *O*-GlcNAc-regulated splicing of OGT and OGA in additional cell types, including a mouse cell line (Supplementary Figure S5).

### O-GlcNAc is a master regulator of detained intron splicing

Because DI-containing transcripts typically represent a small portion of the RNA pool due to their short half-lives (< 1 h), DI changes can be observed at early time points following a perturbation. Changes in other alternative splicing modes require more time to observe, however, because the stable mRNAs produced must be turned over before a change in isoform composition becomes apparent. We sought to answer whether other changes in splicing occur after a longer period of OGT inhibition. Because the average mRNA half-life is about ∼8.5 h, we chose 6 h as our extended time point where we might detect substantive changes not only in DI splicing, but also in other forms of alternative splicing (AS). At 6 h post-treatment with the OGT inhibitor, we observed a global change in DIs, with 3658 out of 4460 DI changes reaching statistical significance (Figure 4A). Strikingly, the vast majority of these (3651, or 99%) decreased in abundance. We randomly selected for validation eight genes with substantial DI decreases during OGT inhibition. RNA-sequencing traces for these genes are provided in (Supplementary Figure S6). We first confirmed by subcellular fractionation that these introns were correctly classified as DIs; as expected, they were enriched in the nucleus (Figure 4C). RT-qPCR showed that DI abundance in all eight genes was substantially reduced in response to OGT inhibition, confirming the global effect on DI splicing (Figure 4D). The global decrease in DIs observed after 6 h of OGT inhibition suggested that O-GlcNAc affects another layer of DI splicing regulation that is more systemic than the highly specific response observed at 2 h. This global response is evidently conserved in other cell types because the DIs for 6 of the 8 validated genes also decreased in abundance in HCT116 colon cancer cells and mouse embryonic fibroblasts when OGT was inhibited (Supplementary Figure S7).

**Figure 4.**
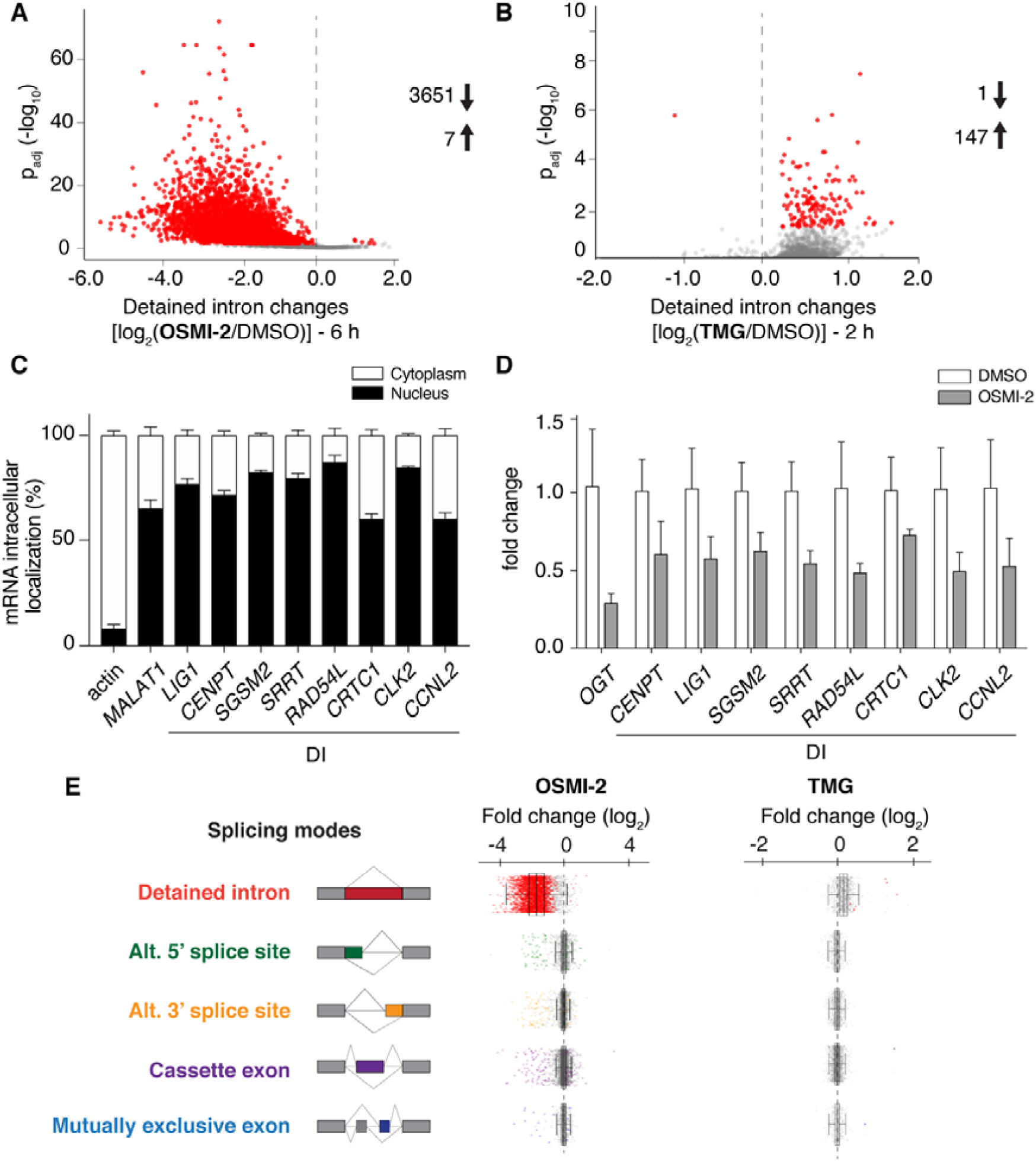
O-GlcNAc regulates DI splicing to control both O-GlcNAc homeostasis and cell state transitions. **(A)** Volcano plot of fold changes (log_2_) in DIs of polyadenylated transcripts isolated from cells treated with OGT inhibitor for 6 h (10 μM OSMI-2 relative to DMSO) showing a global decrease in DI abundance; (**B**) A similar volcano plot as in (A), but generated from cells treated with OGA inhibitor for 2 h (5 uM TMG), showing a global shift toward increased intron detention. **(C)** RT-Qpcr analysis comparing the abundance of DI-containing transcripts in the nuclear and the cytoplasmic fractions for a panel of genes. mRNA was isolated from subcellular fractions obtained from HEK293T cells. The transcript for actin acts as a cytoplasmic localization control, and the transcript for *MALAT1* acts as a nuclear localization control. **(D)** RT-qPCR analysis of the abundance of a panel of DI-containing transcripts from HEK293T cells that were treated with either DMSO or OSMI-2 for 6 h. Transcript abundances are normalized to DMSO control. **(E)** Changes in alternative splicing events after 6 h treatment with OGT inhibitor (10 uM OSMI-2, left) or OGA inhibitor (5 uM TMG, right) represented as boxplots. Color coded dots represent splicing events that meet the statistical cutoff. The black dotted line denotes no change compared to the DMSO control.

We asked whether the global decrease in DIs could be due to decreases in total transcript levels for the affected genes because several findings have suggested that O-GlcNAc levels can affect transcription in a variety of ways. Numerous transcription factors as well as the C-terminus of RNA Pol II are known to be O-GlcNAc modified (50, 67), and OGT has been observed to be enriched at promoters (68). We analyzed changes at the total gene transcript level using RNA-seq data and found that 635 genes showed at least a 2-fold decrease in transcription, while 350 genes showed at least a 2-fold increase, with >0.95 posterior probability of differential expression being considered significant (Supplementary Figure S8). However, among all genes for which total transcript levels decreased, only 34 contained DIs that decreased when OGT was inhibited. We therefore concluded that changes in total transcript quantities cannot account for the effect of reduced O-GlcNAc on DI splicing.

We reasoned that if the global effect we observed on DI levels is dependent on O-GlcNAc, then OGA inhibition should cause an increase in DI abundance. We performed another round of RNA-seq on poly(A)-selected RNA after TMG treatment. At both 2 h and 6 h, we observed a systematic increase in DI abundance that paralleled the systematic decrease observed upon OGA inhibition (Figure 4B and 4E). Fold changes were smaller, and fewer DIs met statistical significance when OGA was inhibited compared to OGT inhibition, possibly because O-GlcNAc levels under steady-state conditions were sufficiently high that the system was resistant to further change. Nevertheless, there was a clear global trend toward increased intron detention as O-GlcNAc levels increased. Taken together, the opposing effects of OGT and OGA inhibition on DI abundance show that cells effect a global DI splicing switch in response to changing O-GlcNAc levels.

The striking impact of O-GlcNAc on DI splicing pathways prompted us to ask whether there were global changes in other forms of alternative splicing. We examined four canonical forms of alternative splicing after OGT inhibition for 6 h, and compared the median fold changes to that of DIs (Figure 4D). DIs displayed a median 3-fold decrease [log_2_(fold-change) of −1.6], but the median fold-changes for other alternative splicing modes were negligible. Moreover, of all non-DI alternative splicing events, only 5% (330 events) reached statistical significance (Supplementary Figure S9). Similar results were observed after OGA inhibition in that the log_2_ median fold change was > 0 for DI splicing, but remained essentially unchanged for other alternative splicing modes. Therefore, the global change in DI splicing observed when O-GlcNAc levels were perturbed was not paralleled by system-wide changes in other forms of alternative splicing. We conclude that O-GlcNAc serves as a global regulator of DI-containing isoform production versus co-transcriptional splicing, thereby linking nutrient levels to mRNA abundance through activity of the *O*-GlcNAc cycling enzymes, particularly OGT.

## DISCUSSION

There are two important findings from this study (Figure 5). First, we found that OGT and OGA levels, which are known to change reciprocally when either enzyme is blocked (50,66), are regulated by detained intron splicing in an O-GlcNAc-dependent manner. These rapid splicing changes serve to maintain O-GlcNAc homeostasis by tuning levels of the O-GlcNAc cycling enzymes as cellular conditions change. The productive versus non-productive forms of the fully-spliced mRNAs are dictated by alternative exon inclusion/skipping for both *OGT* and *OGA*. This allows the switch between DI inclusion and co-transcriptional splicing to be regulated through the same mechanism, even though the alternative exon splicing accomplishes opposite effects on the ratio of the productive and unproductive mRNAs for *OGT* and *OGA*. Second, a reduction in O-GlcNAc that is not rescued by reciprocal, rapid alterations in OGT and OGA causes a global reduction in detained introns, leading to changes in levels of functional mRNA for genes under detained intron splicing control. This global response may serve to reestablish a new setpoint for gene expression under low nutrient conditions. The temporal difference between these two responses suggests that two distinct splicing processes control them. Our recent development of highly specific, fast-acting inhibitors of OGT proved critical for enabling the dissection of these temporal patterns and allowed us to identify direct effects of *O*-GlcNAc on splicing with high confidence.

**Figure 5.**
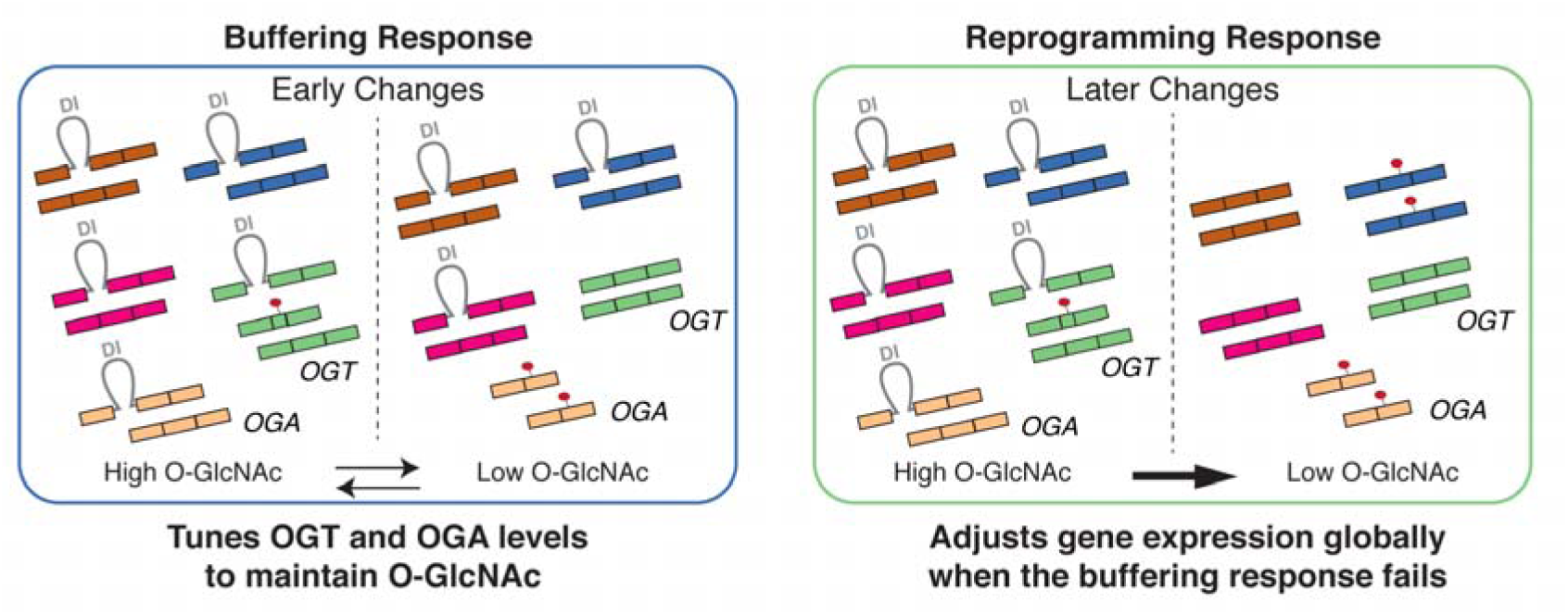
Schematic of O-GlcNAc promoted DI regulation. Models summarizing early and later responses to perturbations in O-GlcNAc levels. Left panel: O-GlcNAc-dependent feedback regulation of DI splicing in *OGT* and *OGA* buffers fluctuations in nutrient availability by tuning levels of productive and non-productive mRNA isoforms to control enzyme abundance. Right panel: Changes in O-GlcNAc levels that exceed the buffering capacity trigger a global DI splicing response to increase (high O-GlcNAc) or decrease (low O-GlcNAc) DI levels to promote an adaptive cell state transition.

The OGT and OGA detained introns contain regions that are extremely conserved across vertebrates, with a 397 nucleotide ‘ultraconserved element’ in OGT that is 100% identical between human, mouse, and rat (69). This degree of conservation is unusual within introns and suggests a conserved mechanism of regulating mRNA levels through DIs in response to changes in O-GlcNAc. This observation highlights the importance of maintaining O-GlcNAc homeostasis even as conditions change. Detained introns are found in other genes involved in metabolism, including *MAT2A*, required for S-adenosylmethionine (SAM) synthesis. The *MAT2A* detained intron is involved in a feedback loop that adjusts *MAT2A* mRNA levels in response to changes in methionine (70). Under low SAM conditions, the RNA methyltransferase METTL16, which uses SAM as a substrate, binds to the *MAT2A* RNA and induces splicing of the *MAT2As* intron. Although molecular details of *MAT2A* DI splicing are still lacking, MAT2A provides the best understood example of how an enzyme can regulate its own DI.

The molecular mechanism underlying O-GlcNAc regulation of DI splicing of *OGT* or *OGA* is still unclear. It is safe to say that it must involve a factor possessing an O-GlcNAc-dependent activity that influences splicing specifically at the *OGT* and *OGA* DIs. It is likely that the extreme nucleotide sequence conservation found within regions of these DIs is required for the specific regulation of the introns. Indeed, many ultraconserved intronic elements occur in RNA binding proteins that auto-regulate the splicing of their own transcripts (71). The structure of *OGT’s* DIs flanking a poison exon is reminiscent of the DIs found in virtually all of the genes encoding the canonical SR proteins (63), and similarly, the structure of *OGA’s* DIs, which flank an essential exon that, when skipped, produces non-functional mRNA, is reminiscent of the DIs found in *Clk1/Clk4* (24). Both the SR proteins and the CLK kinases are regulated via their respective detained introns and contain ultraconservative elements. While SR proteins directly bind their pre-mRNA to affect splicing, the CLK kinases are not known to do so, and so presumably act through their kinase activity to affect the assembly of prespliceosome components, similar to the presumed action of OGT. We attempted to identify factors important for O-GlcNAc-dependent DI splicing of *OGT* by cross-referencing data sets of known O-GlcNAcylated splicing factors with our phosphoproteomic data and then testing whether knockdown of selected factors affected the responsiveness of DI splicing to O-GlcNAc levels (72-74). DI changes in *OGT* were no longer sensitive to O-GlcNAc when core-splicing factors were knocked down, confirming the changes as a splicing-regulated process. However, none of the alternative splicing factors tested as candidates for O-GlcNAc-responsive DI splicing caused a notable change in O-GlcNAc-dependent splicing of *OGT* (Supplementary Figure S10). Given how many cellular components involved in transcription and splicing are modified by OGT, we suspect it will be challenging to unravel the mechanism of O-GlcNAc-dependent splicing regulation. It remains a mystery how changes in O-GlcNAc could accomplish the extreme selectivity observed in the splicing events examined here and how the ultraconserved elements in the genes for the O-GlcNAc cycling enzymes contribute to the regulation of these feedback loops.

The global effect on DI splicing seen at later time points is presumably the result of a different mechanism than the rapid response observed for the *OGT* and *OGA* DIs. One possibility is that a core spliceosome component or peripheral RNA-binding splicing regulator responsible for the DI regulation is O-GlcNAc modified but is inaccessible to rapid turnover through OGA activity. In this case the splicing factor, a number of which are known to be O-GlcNAc modified, may have to undergo protein degradation to turn over the pool of modified protein and thereby affect the splicing of many DIs. This would account for the delay in DI splicing changes between 2h and 6h we observed here. A second possibility is that the modified factor is involved in chromatin modification, nucleosome remodeling, DNA methylation, or the basal transcription machinery, as effectors of all of these pathways are known to be O-GlcNAc modified (5). Again, there could be a lag in the reconfiguration of these factors necessary to either directly affect splicing, or to change the expression of some necessary component. Indeed, differences in O-GlcNAc resistance to removal by OGA have been observed at different loci undergoing chromatin modification (75). A third intriguing possibility comes from recent findings that many RNA-binding factors are involved in liquid-liquid phase transitions that partition RNA and its processing factors into membraneless subcellular compartments (76-78). It is not difficult to imagine how sequestration of DI-containing or NMD-sensitive isoforms within such a compartment could contribute to changes in the stability or processing of transcripts, and that O-GlcNAc could modulate these phase transitions by altering multivalent interaction dynamics between the proteins and RNAs involved (53). The complexity of the O-GlcNAc-modified proteome constitutes the highest barrier to dissecting the exact factors controlling these striking examples of splicing-dependent gene regulation (4,79,80).

## DATA AVAILABILITY

RNAseq data will be deposited in GEO under accession XXXXXXXXX. Custom analysis scripts will be made available upon request.

## SUPPLEMENTARY DATA

Supplementary Data are available at NAR online.

## ACKNOWLEDGEMENT

We thank Phil Sharp, in whose laboratory a portion of the work was performed, for his support and insight. We also thank Dr. Harri Itkonen for the helpful suggestions on the O-GlcNAc pulldown experiments.

## FUNDING

This work was supported by the National Institutes of Health [GM094263 to S.W., BB123456 to J.P., S.G., GM034277 to P.B.]; Singapore A*STAR NSS fellowship to Z.T. University of Rochester Health Sciences Center for Computational Innovation high performance computational resource grant to P.B.; Division of Preclinical Innovation, National Center for Advancing Translational Sciences to D.D., C.J.

Funding for open access charge: National Institutes of Health.

## CONFLICT OF INTEREST

None declared.

