## Supplementary Data for "O-GlcNAc regulates gene expression by controlling detained intron splicing"

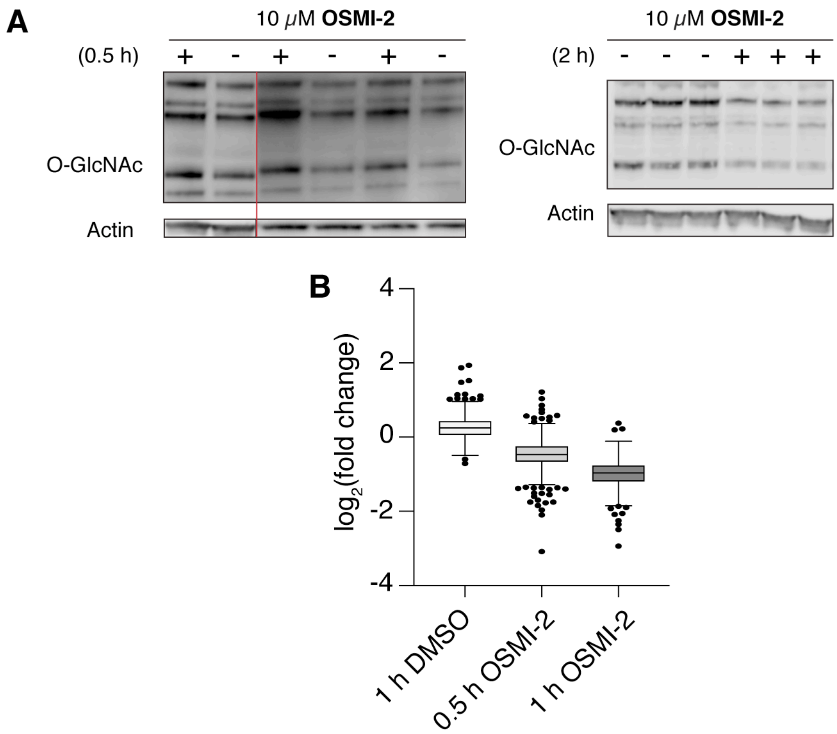


**Supplementary Figure 1. (A)** Western blot for O-GlcNAc levels (α-RL2) after treatment of cells with inhibitors at 10 μM OSMI-2 for 0.5 h (left) or 2.0 h (right). Triplicate lanes for each treatment condition represent biological replicates. **(B)** Box plot of protein abundance, enriched with a combination of RL2 and CTD110.6, after treatment with OSMI-2 for various time point. 1 h DMSO indicates the protein abundance of 1 h DMSO compared to 0.5 h DMSO. DMSO does not decrease global O-GlcNAcylatin.

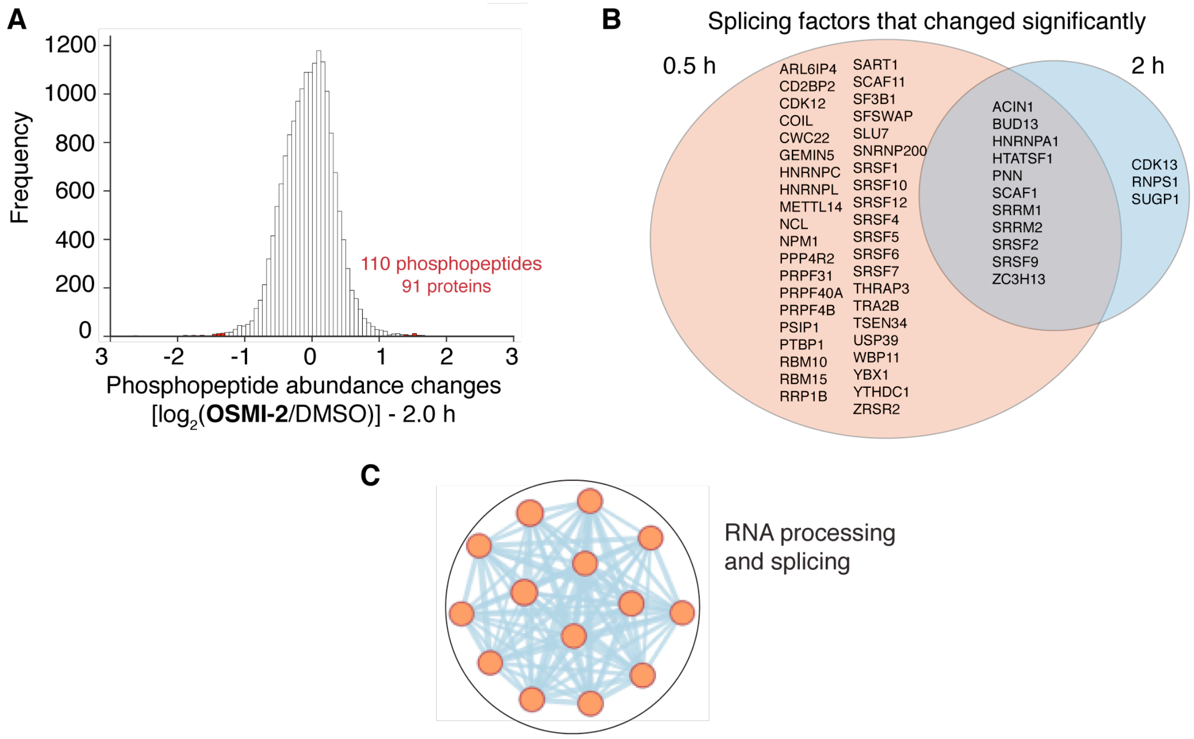


.

**Supplementary Figure 2. (A**) Distribution of phosphopeptide abundances after 2 h treatment with OSMI-2, relative to DMSO control. Hits showed greater than a 2.5-fold change (2 STD from the median; shown in red). **(B)** Comparison of splicing-related genes whose proteins displayed differential phosphorylation (>2.5 fold change, corresponding to 2.0 STD from median) at either 0.5 h or 2 h time points. The list of splicing-related genes were curated mainly using GO. **(C)** Gene Ontology analysis of proteins displaying differential phosphorylation at both 0.5 h and 2 h time points; proteins from the intersection of the Venn diagram in **(B)** was used for analysis. No other module resulted from the analysis.

**A**

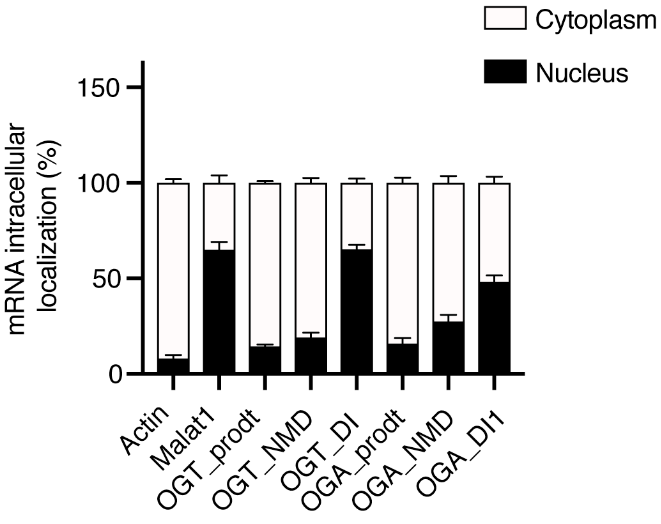


**B**

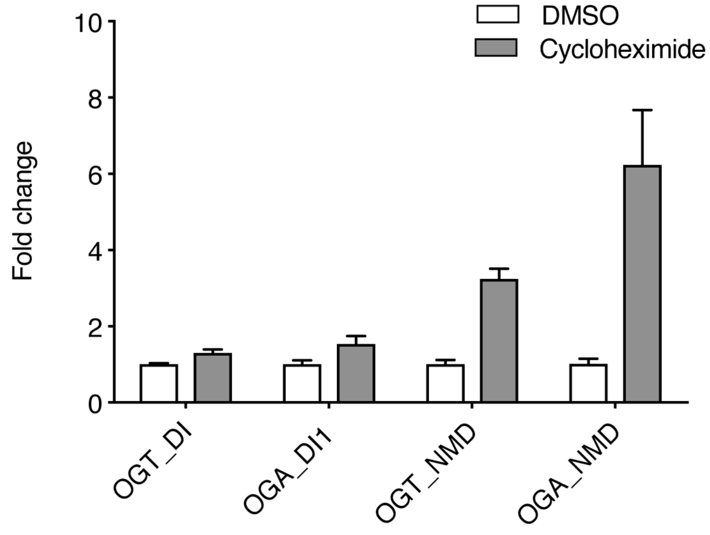


**Supplementary Figure 3. (A)** RT-qPCR analysis comparing the abundance of various *OGT* and *OGA* isoforms in the nuclear and the cytoplasmic fractions. mRNA was isolated from subcellular fractions obtained from HEK293T cells. The transcript for actin acts as a cytoplasmic localization control, and the transcript for *MALAT1* acts as a nuclear localization control. **(B)** RT-qPCR analysis of the abundance of various *OGT* and *OGA* isoforms from HEKs treated 50 µg/mL cycloheximide, a translation inhibitor, for 2 h.

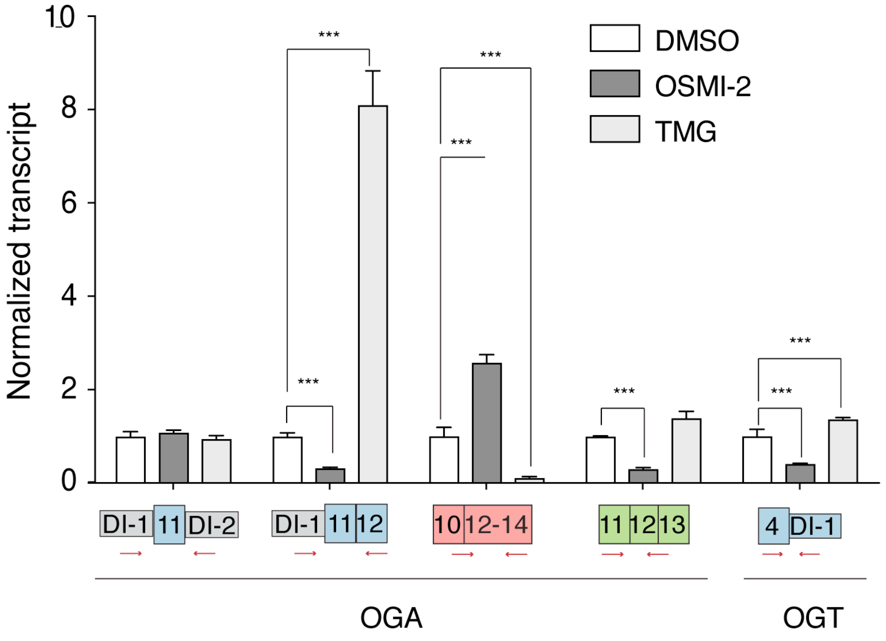


**Supplementary Figure 4.** RT-qPCR results treated for 6 h with either DMSO control (white), OSMI-2 (dark grey), or TMG (light grey). Red arrows indicate binding region for qPCR primers. Levels for each qPCR product were normalized to actin in a given treatment condition, and fold changes were calculated relative to DMSO control. (n ≥ 3 biological replicates; mean ± s.d., *$P \leq 0.05$, ***$P \leq 0.001$, two-tailed Student's *t*-test).

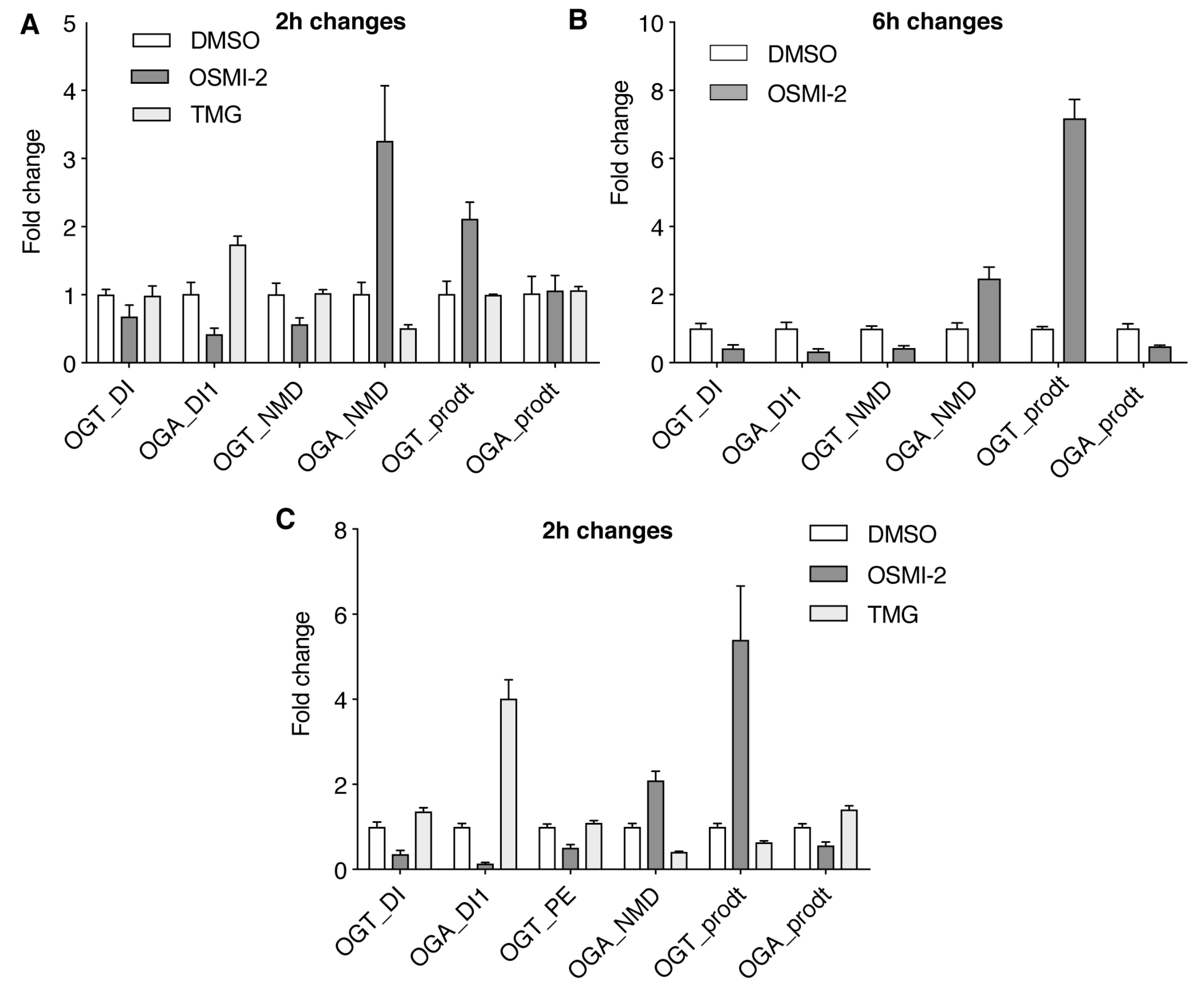


**Supplementary Figure 5.** RT-qPCR analysis of the abundance of various *OGT* and *OGA* isoforms from HCT116 cells treated for 2 h with DMSO, OSMI-2, or TMG **(A)**, or 6 h with DMSO or OSMI-2 **(B)**. Transcript abundances are normalized to DMSO control. **(C)** RT-qPCR analysis of the abundance of various *OGT* and *OGA* isoforms from MEF cells treated for 2 h with DMSO, OSMI-2, or TMG. Levels for each qPCR product were normalized to actin in a given treatment condition, and fold changes were calculated relative to DMSO control. (n ≥ 3 biological replicates; mean ± s.d.)

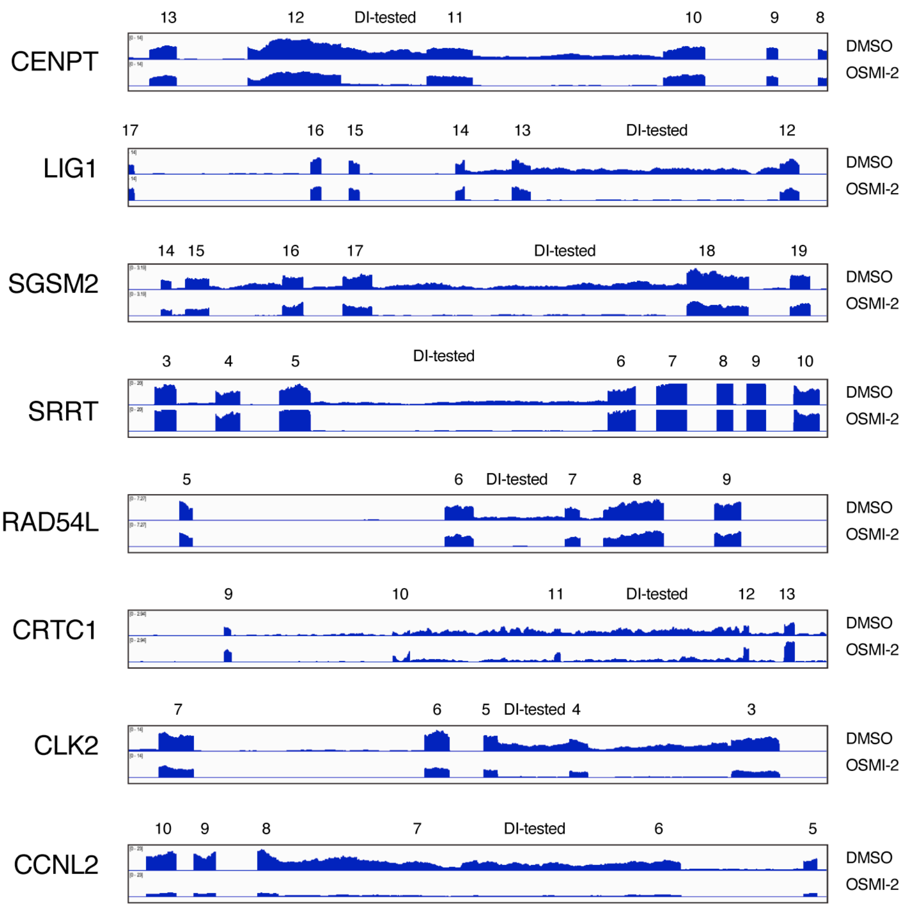


**Supplementary Figure 6.** RNA-seq traces for a panel of DI-containing genes showing the normalized poly(A) read counts in the mRNA isolated from cells treated with either DMSO or OSMI-2.

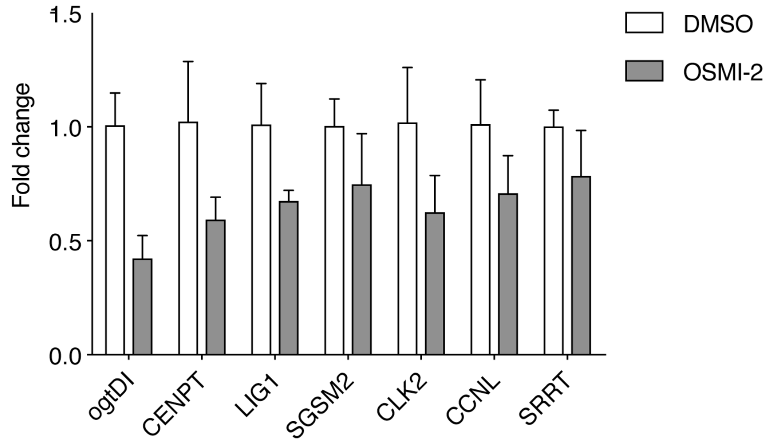


**Supplementary Figure 7.** RT-qPCR results validate DI splicing analysis in HCT116 cells. The panel of genes with DI-containing transcripts were chosen based on those whose DIs displayed changes in abundance in HEK293T cells. cDNA generated from poly(A)-mRNA isolated from either DMSO control or OSMI-2 treated cells for 6 h were used for the RT-qPCR. Levels for each qPCR product were normalized to actin in a given treatment condition, and fold changes are calculated relative to DMSO control. (n ≥ 3 biological replicates; mean ± s.d.)

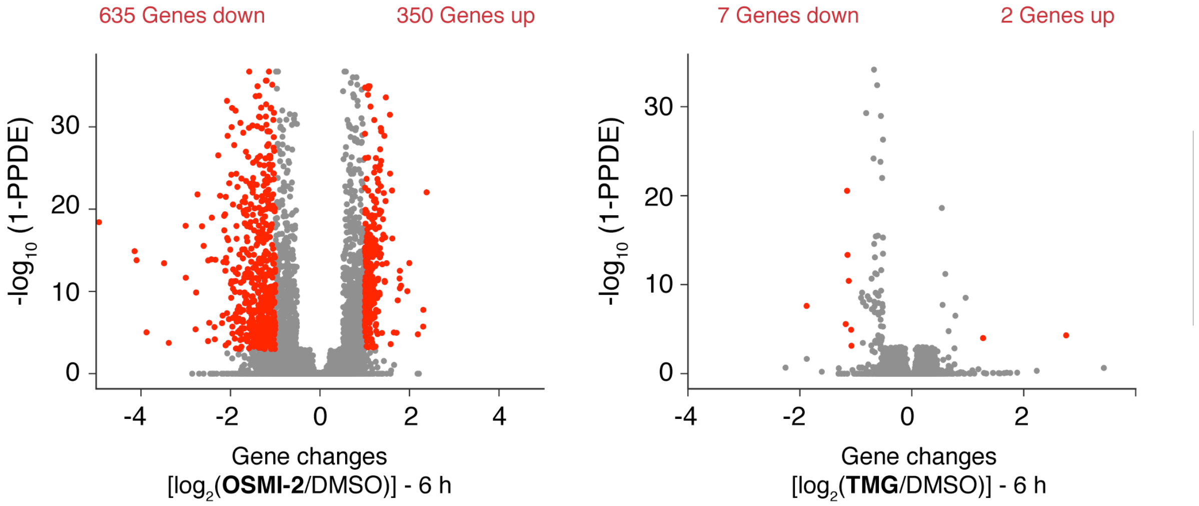


**Supplementary Figure 8.** Volcano plots for gene-level transcript abundances are shown for cells treated with 10 μM OSMI-2 (left) or 5 μM TMG (right). Red dots indicate genes that meet the statistical significance cutoff (PPDE > 0.95) and the fold change cutoff (>2-fold). Of the 635 significantly downregulated genes, only 34 contain a DI.

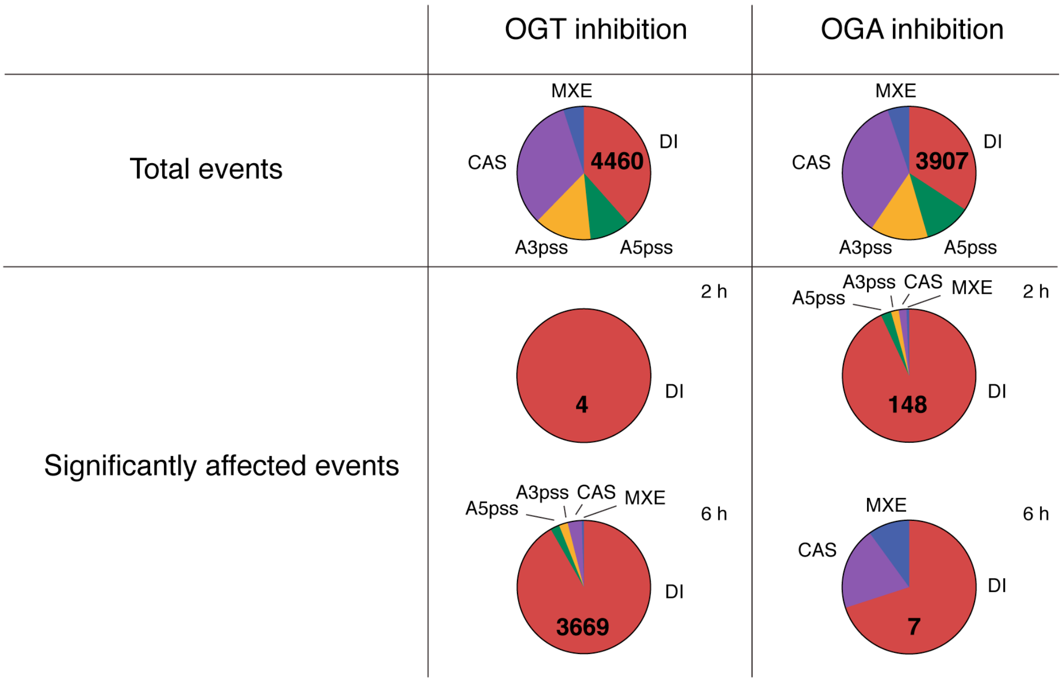


**Supplementary Figure 9.** Comparison of different alternative splicing events following either OSMI-2 or TMG treatment. Total events refer to all canonical alternative splicing events (including DIs) that are detected. Significantly affected events refer to only those that meet the statistical significance cutoff of FDR < 0.05.

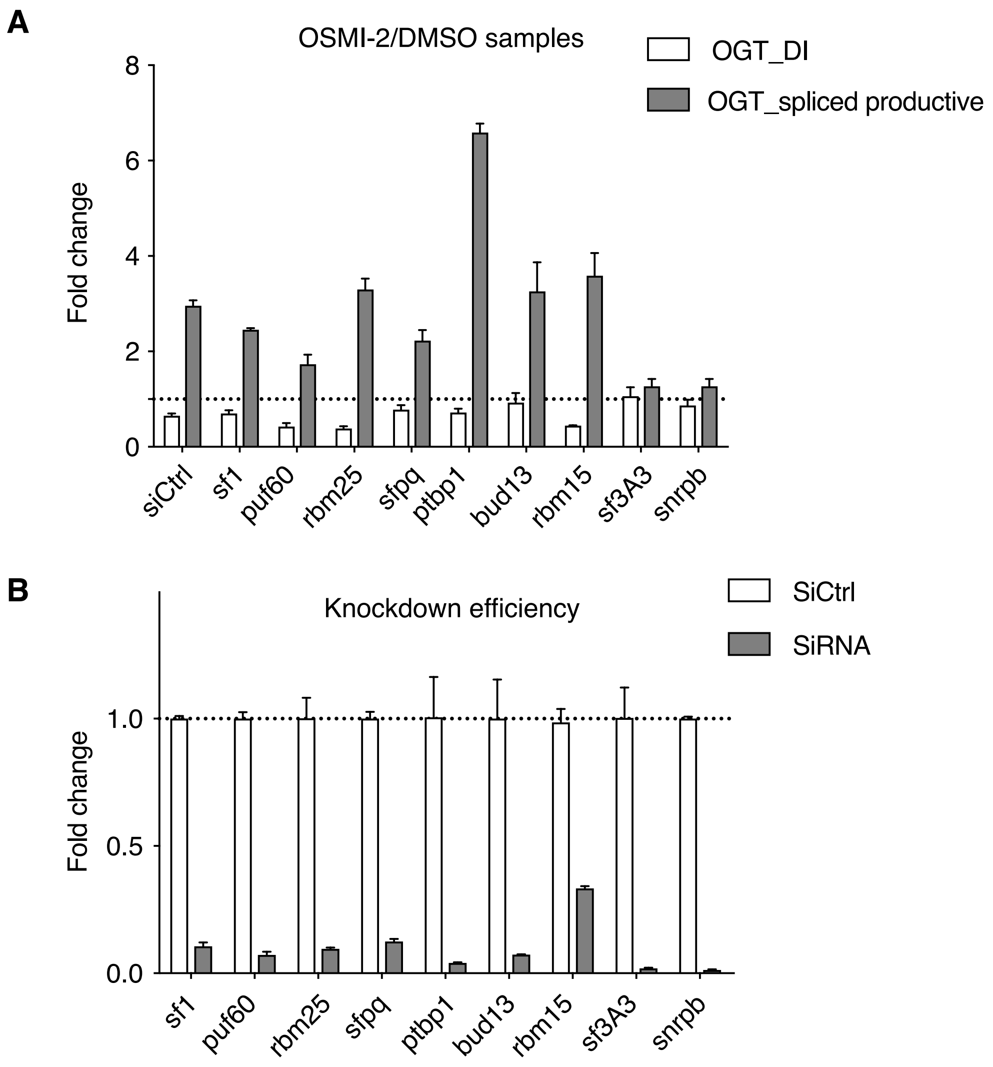


**Supplementary Figure 10. (A)** RT-qPCR analysis of the abundance of *OGT* DI-containing and productive isoforms in HEK293T cells transfected with siRNAs to various putative DI-regulating splicing factors. Cells were treated with either DMSO or OSMI-2 for 2 h, and fold change in isoform abundance in the OSMI-2 sample are calculated relative to DMSO control. SF3A3 and SNRPB are core splicing factors. (n ≥ 2 biological replicates; mean ± s.d.). **(B)** RT-qPCR analysis showing the efficacy of siRNA-mediated knockdown of splicing factors. Knockdown efficiencies are normalized to siCtrl. (n ≥ 2 biological replicates; mean ± s.d.)

| | |
|---|---|
| Primers for OGT | |
| OGT_DI1_FWD<br>OGT_DI1_REV | 5’-ACTGTGTTCGCAGTGACCTG-3’<br>5’-GATTCGTGCCGTCAGTTTTC-3’ |
| OGT_DI2_FWD<br>OGT_DI2_REV | 5’-CCTATGCTGCAGGGTCACTT-3’<br>5’-AAGTTCGGTTGCGTCTCAAT-3’ |
| OGT_PE(NMD)_FWD<br>OGT_PE(NMD)_REV | 5’-ACTGTGTTCGCAGTGACCTG-3’<br>5’-GAGGAGGAGGAGGTTGTGTG-3’ |
| OGT_Prod_FWD<br>OGT_Prod_REV | 5’-ACTGTGTTCGCAGTGACCTG-3’<br>5’-CAAATTTCCCCTTGTGCATT-3’ |
| OGT_DI_MUS_FWD<br>OGT_DI_MUS_REV | 5’-CCTGGGTCGCTTGGAAGAA-3'<br>5’-CCCAGTCCTGCTCTAGTCTG-3' |
| OGT_PE_MUS_FWD<br>OGT_PE_MUS_REV | 5’-CCTGGGTCGCTTGGAAGAA-3'<br>5’-GACTTTTCGGTGCGTTCCTT-3' |
| OGT_Prod_MUS_FWD<br>OGT_Prod_MUS_FWD | 5’-CTGTGTTCGCAGTGACCT-3'<br>5’-CAGCCAAATCTCCCCTTGTG-3' |
| Primers for OGA | |
| OGA_DI1_DI2_FWD<br>OGA_DI1_DI2_REV | 5’-ACCAATCCTTCCAGCCATCT-3’<br>5’-GGACATTTGAGACCCCAGGA-3’ |
| OGA_DI1_FWD<br>OGA_DI1_REV | 5’-TGGAGGTAGGAGTCAGTGGA-3’<br>5’-GGACATTTGAGACCCCAGGA -3’ |
| OGA_NMD_FWD<br>OGA_NMD_REV | 5’-GGAAAGCAGCCCTCCTACTA-3’<br>5’-TTTGTACAGTGGTTAGCGTTTG-3’ |
| OGA_Prod_FWD<br>OGA_Prod_REV | 5’-TCTTGTACACGGATGCCTCA-3’<br>5’-AGAACCCTGGGCCTTTAGAG-3’ |
| OGA_DI_MUS_FWD<br>OGA_DI_MUS_REV | 5’-TCACAGCCTTGCCTTGAAAC-3'<br>5’-TTGGAGGTTGGAGTCAGTGG-3' |
| OGA_NMD_MUS_FWD<br>OGA_NMD_MUS_REV | 5’-GTGCAGTGGTTAGCGTTTG-3'<br>5’-GACGAAGCAGTAATCCAGGC-3' |
| OGA_Prod_MUS_FWD<br>OGA_Prod_MUS_FWD | 5’-TCGTAGTCACTCTTCAGCACA-3'<br>5’-TCTTGTACACGGAAGCCTCA-3' |
| Primers for other DIs | |
| CENPT_DI_FWD<br>CENPT_DI_REV | 5’-TGTCAACTCTCACCCAAGCT-3’<br>5’-GTGTATCACTCCCTGCCCTG-3’ |
| LIG1_DI_FWD<br>LIG1_DI_REV | 5’-CTCCACCATCTCTCCGATCC-3’<br>5’-GGGTGATGGTGTCCTTCTCA-3’ |
| SGSM2_DI_FWD<br>SGSM2_DI_REV | 5’-CCCTTCAGTAGCCACAGGAA-3’<br>3’-GGGGTTCCAGATCATCCACT-3’ |
| SRRT_DI_FWD<br>SRRT_DI_REV | 5’-CTGGAACCCCAAAATAAGCA-3’<br>5’-ATAGTACGCGTCCCCAAACA-3’ |
| RAD54L_DI_FWD<br>RAD54L_DI_REV | 5’-CGCTCATGACCAGCTGAAG-3’<br>5’-ACATGACCACAGGCTCTTCC-3’ |
| CRTC1_DI_FWD<br>CRTC1_DI_REV | 5’-GCTTGAGGAGAGGAGATGGG-3’<br>5’-ATAGTACGCGTCCCCAAACA-3’ |
| CLK2_DI_FWD<br>CLK2_DI_REV | 5’-CATTGTACAACTCGGCCGAA-3’<br>5’-CTTCCCCTGTGACCATCTGT-3’ |
| CCNL2_DI_FWD<br>CCNL2_DI_REV | 5’-GCAAGATAAATGCAGGCACA-3’<br>5’-AGATAATGCAGGCGGAAGAC-3’ |
| Primers for KD targets | |
| SF1_FWD<br>SF1_REV | 5’-GGAGCGGCACAACCTCATC-3’<br>5’-CCGGATCATAATCTTGGCATTGC-3’ |
| PUF60_FWD<br>PUF60_REV | 5’-TCGTGGAGTATGAGGTCCCC-3’<br>5’-CCTGCCCTATGTTGCTGGG-3’ |
| RBM25_FWD<br>RBM25_REV | 5’-TTTCCACCTCATTTGAATCGCC-3’<br>5’-AGTGGGTACTAAGACAGTTGGAG-3’ |
| SFPQ_FWD<br>SFPQ_REV | 5’-AGCGATGTCGGTTGTTTGTTG-3’<br>5’-AGCGAACTCGAAGCTGTCTAC-3’ |
| PTBP1_FWD<br>PTBP1_REV | 5’-AGCGCGTGAAGATCCTGTTC-3’<br>5’-CAGGGGTGAGTTGCCGTAG-3’ |

| | |
|---|---|
| BUD13_FWD<br>BUD13_REV | 5’-CTTTCCAAGGCCGAGTATCTG-3’<br>5’-ATCATCCACAATCCGCATTCC-3’ |
| RBM15_FWD<br>RBM15_REV | 5’-ACGACCCGCAACAATGAAG-3’<br>5’-GGAAGTCGAGTCCTCACCAC-3’ |
| SF3A3_FWD<br>SF3A3_REV | 5’-GTCATGGCTAAAGAGATGCTCACC-3’<br>5’-TCCTCCTTTCGTAATCCATCCTT-3’ |
| SNRPB_FWD<br>SNRPB_REV | 5’-CTGGTCTCAATGACAGTAGAGGG-3’<br>5’-GGGACCCATAGGAGGTCTCATA-3’ |
| Primers for control transcripts | |
| MALAT1_FWD<br>MALAT1_REV | 5’-CCTGCAAATTGTTAACAGAA-3’<br>5’-TCAGCTTCCGCTAAGATGCTAGCTT-3’ |
| ACTB_FWD<br>ACTB_REV | 5’-TGGGACGACATGGAGAAAAT-3’<br>5’-AGAGGCGTACAGGGATAGCA-3’ |
| ACTB_MUS_FWD<br>ACTB_MUS_REV | 5’-GGCTGTATTCCCCTCCATCG-3'<br>5’-CCAGTTGGTAACAATGCCATGT-3' |

**Supplementary Table 1.** Primers used for quantitative real-time PCR for OGT and OGA. Primers for experiments done in human cell lines were designed based on Hg19 human complete genome sequence. Primers for experiments done in mouse cell lines were designed based on GRCm38 complete mouse genome sequence.

| siRNAs | |
|---|---|
| siCtrl | Silencer Negative Control No.1 siRNA |
| siSF1 | cat. s224783 |
| siPUF60 | cat. s223538 |
| siRBM25 | cat. s33914 |
| siSFPQ | cat. s12712 |
| siPTBP1 | cat. s11436 |
| siBUD13 | cat. s39455 |
| siRBM15 | cat. s34936 |
| siSF3A3 | cat. s21534 |
| siSNRPB | cat. s13219 |

**Supplementary Table 2.** siRNAs used for knockdown experiments and their catalogue numbers. All siRNAs were obtained from Thermo Fisher.
